# Detection of antibodies neutralizing historical and emerging SARS-CoV-2 strains using a thermodynamically coupled de novo biosensor system

**DOI:** 10.1101/2021.06.22.449355

**Authors:** Jason Z. Zhang, Hsien-Wei Yeh, Alexandra C. Walls, Basile I.M. Wicky, Kaiti Sprouse, Laura A. VanBlargan, Rebecca Treger, Alfredo Quijano-Rubio, Minh N. Pham, John C. Kraft, Ian C. Haydon, Wei Yang, Michelle DeWitt, Cameron Chow, Lauren Carter, Mark H. Wener, Lance Stewart, David Veesler, Michael S. Diamond, David Baker

## Abstract

With global vaccination efforts against SARS-CoV-2 underway, there is a need for rapid quantification methods for neutralizing antibodies elicited by vaccination and characterization of their strain dependence. Here, we describe a designed protein biosensor that enables sensitive and rapid detection of neutralizing antibodies against wild type and variant SARS-CoV-2 in serum samples. More generally, our thermodynamic coupling approach can better distinguish sample to sample differences in analyte binding affinity and abundance than traditional competition based assays.

## Main

With the availability of COVID-19 vaccines, the rise of more transmissible and pathogenic virus mutants^1^, and known time-dependent declines in immunity following infection^2^, there is a need to determine the degree of serological antibody protection from SARS-CoV-2. Knowledge of individual immunity to SARS-CoV-2 is useful not only to determine personal actions, but also to guide early therapy of patients and evaluate the efficacy of antibody treatment and vaccines over time against emerging viral variants of concern (VoC)^3,4^.

The receptor binding domain (RBD) of the SARS-CoV-2 spike protein binds to the angiotensin-converting enzyme 2 (ACE-2) receptor on target cells and is the immunodominant target of neutralizing antibodies (nAbs) identified from convalescent and post-vaccination plasma^5^. Several SARS-CoV-2 VoCs have exploited this and acquired mutations in the RBD, which allow for escape from nAbs^3,4,6^ targeting WT. Serological antibody tests, ideally home-based diagnostics, are critical to evaluate the response to vaccination and viral infection^2^. Assays that measure antibody titer and neutralizing capability exist but are not compatible with home use. Traditional affinity-based immunoassays, such as ELISA assays^7^, can quantitatively measure antibody titer but due to inherent complexity and instrumentation they require a centralized laboratory for diagnostics. Antibody neutralizing capabilities are traditionally measured in cell-based live viral infection assays that require BSL3 facilities^8^. Lateral flow antigen tests^9^ have been introduced but they are used primarily as binary qualitative tests and report only binding between antibody-antigen rather than neutralization^10^. Recently developed cell-free tools can measure antibody titer, but cannot necessarily evaluate neutralization, and none of the currently available tools have estimated neutralization activity against the emerging set of SARS-COV-2 VoCs^11–13^. We aimed to develop a sensor technology that can quantitatively measure nAb responses against different strains of SARS-CoV-2, be adapted for all-in-solution, multi-well format, and provide rapid results in 30 minutes (faster than established ELISA assays measuring SARS-CoV-2 antibody titer (~6 hours) or cell-based neutralization assays(~several days)).

To achieve this goal, we focused on antibodies that interfere with RBD:ACE-2 interactions as a proxy for antibody neutralization^12,13^ (**Fig 1a**). We adapted a designed coronavirus spike RBD biosensor^14^ consisting of a switchable lucCageRBD protein containing a “cage” domain that in the closed state of the sensor binds a “latch” domain containing the picomolar affinity RBD binding LCB1 protein, and a lucKey protein that binds to the open state of the sensor, reconstituting luciferase activity^14^. In the absence of RBD, the sensor is in the closed state with the latch bound to the cage, blocking luciferase reconstitution. Upon addition of RBD, the free energy of binding to lucCageRBD, together with that of lucKey, drives switch opening and generation of luminescent signal (**Fig 1b**). Since the biosensor is under thermodynamic control and fully reversible, it is capable of detecting RBD targeted SARS-CoV-2 antibodies that compete with LCB1 at or near the ACE-2 binding interface of RBD. Starting from the open luminescent state of the sensor bound to the RBD, addition of antibody pulls the equilibrium towards the dark closed state (**Fig 1b**).

**Fig 1:**
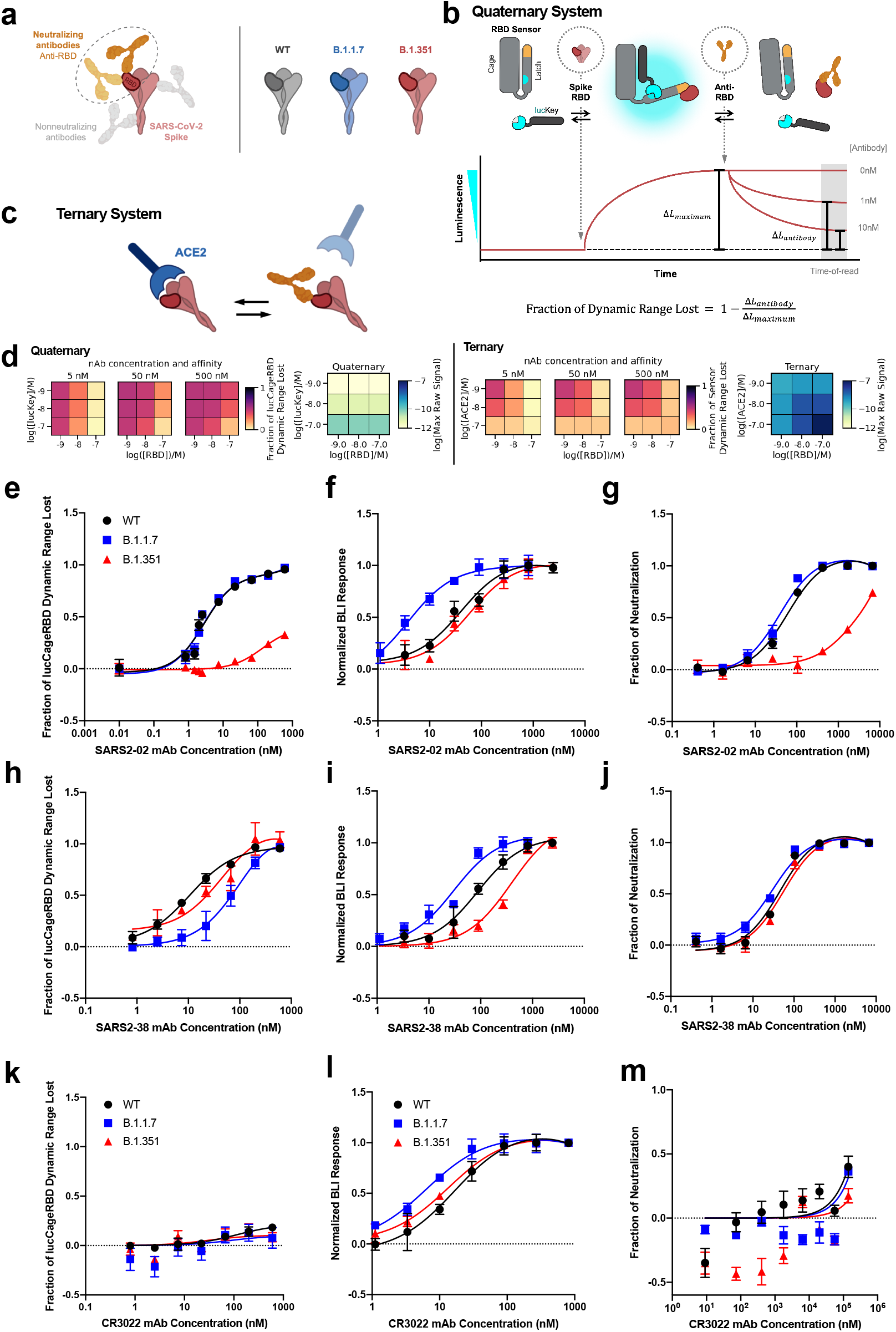
Design and characterization of sensors for monoclonal antibody detection. **a**, To detect neutralizing antibodies which primarily block the interaction between ACE-2 and the receptor binding domain (RBD) of SARS-CoV-2 spike WT and other emerging variants, we designed our lucCageRBD assay, which utilizes the RBD sensor and lucKey. **b-c,** Schematic of the LOCKR nAb biosensor system (quaternary, this work) (**b**) and ACE2:RBD out-competition format (ternary, previous work) (**c**). The RBD sensor contains 2 domains that interact, the Cage and Latch, the latter of which contains smBiT of luciferase (blue) and the *de novo* LCB1 domain (yellow) designed to recognize the ACE-2 binding region of RBD^15^. LucKey contains the Cage-associating key domain and lgBiT of luciferase (blue). RBD WT or variants bind to LCB1, which together with Key:Cage binding enables reconstitution of luciferase, thus increasing luminescence. Neutralizing antibodies compete for RBD binding, thus shifting for Cage:Latch binding, limiting Key:Cage interaction, and disturbing luciferase reconstitution, thus decreasing luminescence. As increasing nAb concentrations should promote decreases in luminosity, we created the fraction of lucCageRBD dynamic range lost metric. **d,** Simulations for the detection and deconvolution of nAb titer from affinity. Each sub-plot corresponds to the sensor response for each setting of the decision matrix, which is defined by a combination of lucKey and RBD concentrations (quaternary system; left) or ACE2 and RBD concentrations (ternary system; right). For each sensor system, the raw maximum signal (absence of nAb, used for signal normalization) is also shown (blue heatmaps). **e-m**, Different concentrations of either SARS2-02 (**e-g**), SARS2-38 (**h-j**), CR3022 (**k-m**) mAbs with 5 nM RBD WT, B.1.1.7, and B. 1.351 were tested in the lucCageRBD assay (**e, h, k**), BLI for binding to RBD strains (**f, i, l**), and spike VoC-containing SARS-CoV-2 live virus (**g, j**) or VSV-based pseudovirus infection (**m**). lucCageRBD and BLI experiments were performed in triplicates, viral infection experiments were performed in duplicate and data are mean ± s.e.m.

Unlike previously described competition assays which directly assess the extent of ACE-2:RBD complex formation (by luciferase reconstitution^12^ or capturing of enzyme conjugated to one component^13^) (**Fig 1c**), in this thermodynamic coupling scheme, the RBD is unmodified and free in solution. This simplifies testing reactivity against RBD variants of concern, since no labeling is required. A more fundamental advantage, as illustrated by the simulations in **Fig 1d and Extended Data Fig 1**, is that compared to ternary sensor systems that rely on direct competition of the ACE-2:RBD interaction by neutralizing antibodies, our quaternary sensor system can more readily distinguish analyte binding affinity and abundance^17^, both of which are relevant for diagnosing the success of vaccination. Distinguishing signals due to higher concentrations of more weakly binding analytes from those due to lower concentrations of more strongly binding analytes is challenging for a simple competition mechanism because the extent of competition depends on the ratio of competitor concentration to the *Kd.* As shown in **Fig 1d**, decision matrices generated by taking measurements for different configurations of the quaternary sensor system (changing the concentrations of the RBD and lucKey sensor components) enable better discrimination of analyte concentration and affinity across the constant [concentration]/Kd range than measurements for different configurations of the ternary sensor (sensor activation across the full analyte affinity and concentration ranges are shown in **Extended Data Fig 1**). A further advantage of our quaternary system is that the maximum sensor response in the absence of nAb (used for signal normalization; decision matrices at the far right in **Fig 1d**), varies less between the different sensor configurations than in the ternary system because the lucCageRBD concentration is constant and the RBD is unlabeled; variation in maximum signal is a problem for luminescent (or other enzyme coupled) sensors because of substrate depletion effects and instrument detection limits. This variation in maximum signal can be avoided in the ternary system if one component is kept fixed, however in this case the high affinity/low concentration and low affinity/high concentration regimes are even harder to resolve (**Extended Data Fig 1g-h**; substituting ACE-2 for a higher affinity RBD binder like LCB1 does not alleviate this problem). Finally, our quaternary system is considerably more tunable. The affinities of the Latch and Key for the Cage can both be tuned to maximize the dynamic range of the system for the relevant analyte affinities and concentrations, whereas to tune response in the ternary system, possibly deleterious mutations must be made at the interface between interacting partners.

To characterize the quaternary sensor system experimentally, we investigated the modulation of lucCageRBD signal by combinations of RBD (and RBD variants) and RBD binding proteins. Addition of a 333 pM RBD to the sensor resulted in a rapid (t_1/2_=22 min) five fold increase in luminescence from baseline which was rapidly quenched (t_1/2_=10 min) by subsequent addition of 200 nM LCB1, which competitively inhibits RBD binding to the RBD sensor (**Extended Data Fig 2**). lucCageRBD also responds to WT, B.1.1.7, and B.1.351 RBD with different dynamic ranges (DR) and EC50 values (**Extended Data Fig 3**). To quantify the perturbation of the system elicited by a RBD binding protein, we use as a metric the fraction decrease in total DR (**Fig 1b;** this is also used in the simulations above). lucCageRBD dose response curves using the neutralizing monoclonal antibody (mAb) CV30^18^ show that increasing RBD WT concentration increases the lucCageRBD EC50 while increasing sensor concentration (1:1 stoichiometry) decreases the lucCageRBD EC50 (**Extended Data Fig 4**), illustrating how changing RBD and sensor concentrations can tune the sensitivity of the lucCageRBD assay.

To evaluate the detection of nAbs through equilibrium perturbation of the lucCageRBD -- RBD system, we compared binding to the spike, virus neutralization, and sensor activation for a set of five anti-spike mAbs (SARS2-02, SARS2-38, CV30, B38 and CR3022) for the WT, B.1.1.7, and B.1.351 spike variants (**Fig 1e-m and Extended Data Fig 5**). Over this set of antibodies and spike variants, virus neutralization correlates with sensor activation better than with spike binding affinity (**Extended Data Fig 5g-i)**, as expected since the sensor is only sensitive to binding near the ACE-2 binding site which is the target of most neutralizing antibodies. As an example, the SARS2-02 antibody binds (by biolayer interferometry) B.1.351 and WT RBD (**Fig 1f**) with roughly equal affinities, but neutralizes infectious SARS-CoV-2 (live virus) containing WT and B.1.1.7 spike proteins much more potently (20-40-fold increase in IC50) than B.1.351 spike-containing virus^19^ (**Fig 1g**). Consistent with the neutralization results, the SARS2-02 antibody produces a large decrease in lucCageRBD signal with WT and B1.1.7 RBD but a partial response with B.1.351 RBD (EC50 ~40-fold increased) (**Fig 1e, Table 1**). To confirm the ability to differentiate nAb concentration and affinity suggested by the simulations in Fig 1c, we assayed two antibodies (CV30 and B38) with different affinities for WT RBD at two different concentrations each, using four different sensor settings, and found that the differential sensor readings for each condition were consistent with the computational model (**Extended Data Fig. 1g-h**).

**Table 1.**

|  | lucCage RBD |  |  |  |  |  | BLI - binding |  |  | Neutralization |  |  |  |  |  |
| --- | --- | --- | --- | --- | --- | --- | --- | --- | --- | --- | --- | --- | --- | --- | --- |
|  | % DR lost | EC50 (nM) | % DR lost | EC50 (nM) | % DR lost | EC50 (nM) | Kd (nM) | Kd (nM) | Kd (nM) | % neutralization | IC50 (nM) | % neutralization | IC50 (nM) | % neutralization | IC50 (nM) |
| Antibodies | WT RBD | WT RBD | B.1.1.7 RBD | B.1.1.7 RBD | B.1.351 RBD | B.1.351 RBD | WT RBD | B.1.1.7 RBD | B.1.351 RBD | WT RBD | WT RBD | B.1.1.7 RBD | B.1.1.7 RBD | B.1.351 RBD | B.1.351 RBD |
| SARS2-02 | 95.6 +/- 0.133 | 2.82 | 97 +/- 0.215 | 2.8 | 32.8 +/- 0.395 | >600 | 41 | 3.88 | 59.9 | 100 +/- 0 | 60.6 | 100 +/- 0 | 39 | 74.2 +/- 0.9 | 1483 |
| SARS2-38 | 91.4 +/- 0.533 | 10.6 | 89.5 +/- 0.97 | 70.6 | 23.8 +/- 1.35 | 47.6 | 88.1 | 30.3 | 394 | 99.8 +/- 0.215 | 45.7 | 99.7 +/- 0.3 | 30.2 | 100 +/- 0 | 54.9 |
| CV30 | 88.8 +/- 8.16 | 38.6 | 93.5 +/- 4.78 | 25.3 | 94.4 +/- 6.5 | 35.4 | 25.2 | 28.5 | 175 | 89.9 +/- 2.51 | 59.6 | 95.5 +/- 0.984 | 6.76 | -17 +/- 24.4 | >1024 |
| B38 | 81.5 +/- 7.47 | 70.5 | 87.6 +/- 6.95 | 23 | 104 +/- 16.6 | 35.4 | 192 | 148 | 376 | 100 +/- 0 | 362 | 6.7 +/- 22.6 | >7749 | -18.9 +/- 28.4 | >7749 |
| CR3022 | 18.3 +/- .88 | >600 | 7.72 +/- 10.3 | >600 | 8.35 +/- 4.58 | >600 | 17 | 6.42 | 13.6 | 40.1 +/- 8.3 | 175988 | 36.4 +/- 1.28 | >140000 | 17.5 +/- 5.56 | >140000 |

We next investigated whether the correlation between sensor response and neutralizing activity observed over the panel of monoclonal antibodies held for polyclonal antibodies in serum. We first explored whether lucCageRBD could measure SARS-CoV-2 protective antibodies from serum samples from mice vaccinated with spike or SARS-CoV-2-RBD nanoparticles^21^ (**Fig 2a**). As complex biological matrices can affect absolute luminescent readings, we used Antares2 as a BRET reference for internal calibration, and used as a measure of sensor activation the ratio of luminescence signal to the internal standard^14,22^ (see Methods). Prior to vaccination, mouse serum samples did not decrease activation, whereas serum samples post-prime dosing (week 3) and post-boost dosing (week 5) showed progressively larger decreases in activation (**Fig 2a**). Over several mouse serum samples from vaccinated animals, decreases in the luminescence ratio from lucCageRBD correlate (R^2^=0.711) with the log_10_ IC50 values against WT spike-presenting pseudovirus (**Fig 2a**) and the log_10_ reciprocal EC50 titer^23^ measured in ELISA (R^2^=0.917) (**Extended Data Fig 6a**). We next tested lucCageRBD’s utility in serum samples from humans vaccinated with BNT162b2 against WT, B.1.1.7, and B.1.351 RBD^24^. The lucCageRBD loss in DR correlates well with the SARS-CoV-2 (WT) antibody titer detected from ELISA using the log_10_ of the Z-score metric (**Fig 2b and Extended Data Fig 6b and 7**) (R^2^=0.942) and with log_10_ IC50 values against pseudovirus displaying either WT (R^2^=0.832), B.1.1.7 (R^2^=0.89), or B.1.351 spike (R^2^=0.961) proteins (**Fig 2b**).

**Fig 2:**
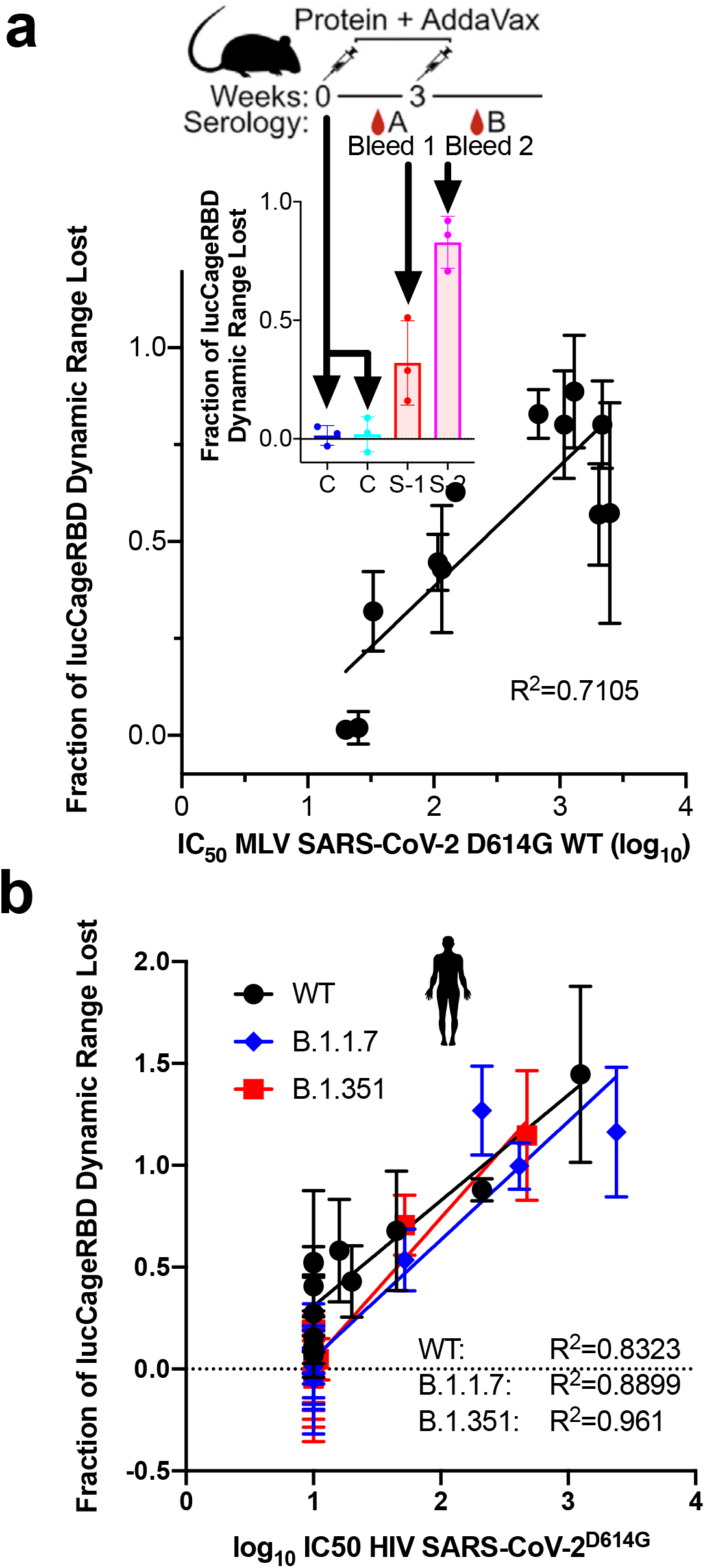
Detection of neutralizing antibodies in vaccinated serum. **a,** Serum (10%) from vaccinated mice were collected and tested for pseudovirus infection and in the lucCageRBD assay (n=12 serum samples). (Inset) Mouse serum samples pre-immunization, post-prime dosing (week 3), and post-boost dosing (week 5) were collected and tested in the lucCageRBD assay. **b,** Serum (10%) from vaccinated patients were tested for pseudovirus infection and in the lucCageRBD assay using RBD WT, B.1.1.7, and B.1.351 (n=12 serum samples). All neutralization experiments were performed in at least duplicates, all lucCageRBD experiments were performed in triplicate and data are mean ± s.e.m.

Our sensor complements previously described COVID-19 serological tests. First, it does not require labeling of the RBD or variant RBDs, which makes it straightforward to substitute in new escape variant RBDs as they are identified. Previous studies have demonstrated detection of antibodies against the WT RBD, here we demonstrate differentiation between antibodies based on their extent of reaction with WT and escape variant RBDs. Second, the components of the sensor can be readily made in *E. coli* and can be lyophilized without loss of performance (**Extended Data Fig 8 and 9**); hence there are potential advantages in shelf life and manufacturing. The low stability of the ACE-2 protein has complicated high throughput, one-step serological detection of nAbs^16^, use of the hyperstable LCB1 instead avoids this problem. Lastly, our thermodynamic coupling based readout enables quantification of both antibody abundance and affinity, which are both relevant for assessing responses to vaccination. Further research will focus on incorporating the sensor into a scalable 384-well high throughput format or a low-cost point-of-care diagnostic testing platform. More generally, with the considerable recent advances that now enable computational design of high affinity binding proteins and the embedding of them in designed protein switches, the approach described in this paper should be readily extensible to quantification of the binding affinity and abundance of a wide variety of analytes of interest.

## Methods

### Cells

HEK293T/17 is a human embryonic kidney cell line (ATCC, CRL-11268). The HEK-ACE2 adherent cell line was obtained through BEI Resources (NR-52511). All adherent cells were cultured at 37°C with 8% CO2 in flasks with DMEM supplemented with 10% FBS (Hyclone) and 1% penicillin-streptomycin.

HEK293F is a female human embryonic kidney cell line transformed and adapted to grow in suspension (Life Technologies). HEK293F cells were grown in 293FreeStyle expression medium (Life Technologies), cultured at 37C with 8% CO2 and shaking at 130 rpm. Expi293F cells are derived from the HEK293F cell line (Life Technologies). Expi293F cells were grown in Expi293 Expression Medium (Life Technologies), cultured at 36.5C with 8% CO2 and shaking at 150 rpm.

Vero E6 (CRL-1586, American Type Culture Collection), Vero-TMPRSS2 (a gift of S. Ding, Washington University) and Vero-hACE2-TMPRSS2 (a gift of A. Creanga and B. Graham, National Institutes of Health (NIH)) cells were cultured at 37°C in Dulbecco’s modified Eagle medium (DMEM) supplemented with 10% fetal bovine serum (FBS), 10 mM HEPES (pH 7.3), 1 mM sodium pyruvate, 1× nonessential amino acids and 100 U mL^-1^of penicillin-streptomycin. Vero-TMPRSS2 cell cultures were supplemented with 5 μg mL^-1^of blasticidin. TMPRSS2 expression was confirmed using an anti-V5 antibody (Thermo Fisher Scientific, 2F11F7) or anti-TMPRSS2 mAb (Abnova, Clone 2F4) and APC-conjugated goat anti-mouse IgG (BioLegend, 405308). Vero-hACE2-TMPRSS2 cell cultures were supplemented with 10 μg ml^-1^of puromycin.

### Monoclonal antibodies

The murine mAbs SARS2-02 and SARS2-38 studied were isolated from BALB/c mice immunized with SARS-CoV-2 spike and RBD proteins and have been described previously^25^. Genes encoding CR3022^26^, B38^27^, and CV30^18^ heavy and light chains were ordered from GenScript and cloned into pCMV/R. Antibodies were expressed by transient co-transfection of both heavy and light chain plasmids in Expi293F cells using PEI MAX (Polyscience) transfection reagent. Cell supernatants were harvested and clarified after 3 or 6 days and protein was purified using protein A chromatography (Cytiva).

### Live virus production

The 2019n-CoV/USA_WA1/2020 isolate of SARS-CoV-2 was obtained from the US Centers for Disease Control. The B.1.1.7 isolate was obtained from a nasopharyngeal sample after propagation on Vero-hACE2-TMPRSS2 cells^3^. The chimeric WA1/2020 displaying B.1.351 virus has been described previously^3^. All viruses were deep-sequenced and tittered on Vero-TMPRSS2 cells, and experiments were performed in an approved Biosafety level 3 facility.

### Pseudovirus production

MLV-based and HIV SARS-CoV-2 S pseudotypes were prepared as previously described^28–30^. Briefly for MLV, HEK293T cells were co-transfected using Lipofectamine 2000 (Life Technologies) with an S-encoding plasmid, an MLV Gag-Pol packaging construct, and the MLV transfer vector encoding a luciferase reporter according to the manufacturer’s instructions. For HIV, HEK293T cells were cotransfected using Lipofectamine 2000 (Life Technologies) with an S-encoding plasmid, an HIV Gag-Pol, Tat, Rev1 B packaging construct, and the HIV transfer vector encoding a luciferase reporter according to the manufacturer’s instructions. Cells were washed 3× with Opti-MEM prior to transfection and incubated for ~5 h at 37°C with transfection medium. DMEM containing 10% FBS was added for ~60 h. The supernatants were harvested by spinning at 2,500*g*, filtered through a 0.45 mm filter, concentrated with a 100 kDa membrane for 10 min at 2,500*g* and then aliquoted and stored at −80°C prior to use.

D614G SARS-CoV-2 S (YP 009724390.1), B.1.351 S and B.1.1.7 S pseudotypes VSV viruses were prepared as described previously^31,32^. Briefly, 293T cells in DMEM supplemented with 10% FBS, 1% PenStrep seeded in 10-cm dishes were transfected with the plasmid encoding for the corresponding S glycoprotein using lipofectamine 2000 (Life Technologies) following manufacturer’s indications. One day post-transfection, cells were infected with VSV(G*ΔG-luciferase) and after 2 h were washed five times with DMEM before adding medium supplemented with anti-VSV-G antibody (I1-mouse hybridoma supernatant, CRL-2700, ATCC). Virus pseudotypes were harvested 18-24 h post-inoculation, clarified by centrifugation at 2,500 x g for 5 min, filtered through a 0.45 μm cut off membrane, concentrated 10 times with a 30 kDa cut off membrane, aliquoted and stored at −80°C.

### Mouse serum

Female BALB/c (Stock: 000651) mice were purchased at the age of four weeks from The Jackson Laboratory, Bar Harbor, Maine, and maintained at the Comparative Medicine Facility at the University of Washington, Seattle, WA, accredited by the American Association for the Accreditation of Laboratory Animal Care International (AAALAC). At six weeks of age, mice were immunized, and three weeks later animals were boosted. Prior to inoculation, immunogen suspensions were gently mixed 1:1 vol/vol with AddaVax adjuvant (Invivogen, San Diego, CA) to reach a final concentration of 0.009 or 0.05 mg/mL antigen. Mice were injected intramuscularly into the gastrocnemius muscle of each hind leg using a 27-gauge needle (BD, San Diego, CA) with 50 mL per injection site (100 mL total) of immunogen under isoflurane anesthesia. To obtain sera all mice were bled two weeks after prime and boost immunizations. Blood was collected via submental venous puncture and rested in 1.5 mL plastic Eppendorf tubes at room temperature for 30 min to allow for coagulation. Serum was separated from hematocrit via centrifugation at 2,000*g* for 10 min. Complement factors and pathogens in isolated serum were heat-inactivated via incubation at 56°C for 60 min. Serum was stored at 4°C or −80°C until use. Mouse sera in this study were used in a previous study^21^.

### Human serum and ELISA

Serial human sera specimens were obtained following immunization of laboratory volunteers with the BNT162b2 mRNA vaccine in an IRB-compliant protocol. Antibodies to the SARS-CoV-2 spike protein were measured using an ELISA specific for anti-S1 IgG (Euroimmun, Mountain Lakes, NJ). Antibody levels were quantified by conversion of the optical density to a z-score relative to pre-pandemic serum anti-S1 IgG concentrations, as previously described^23,33^. Sera tests were serial specimens obtained following immunization of laboratory volunteers in an IRB-compliant protocol.

### General procedures for bacterial protein production and purification

The *E. coli* Lemo21(DE3) strain (NEB) was transformed with a pET29b+ plasmid encoding the synthesized gene of interest. Cells were grown for 24 h in LB medium supplemented with kanamycin. Cells were inoculated at a 1:50 mL ratio in the Studier TBM-5052 autoinduction medium supplemented with kanamycin, grown at 37°C for 2-4 h, and then grown at 18°C for an additional 18 h. Cells were collected by centrifugation at 4,000*g* at 4°C for 15 min and resuspended in 30 mL lysis buffer (20 mM Tris-HCl pH 8.0, 300 mM NaCl, 30 mM imidazole, 1 mM PMSF, 0.02 mg mL^-1^ DNase). Cell resuspensions were lysed by sonication for 2.5 min (5 s cycles). Lysates were clarified by centrifugation at 24,000*g* at 4°C for 20 min and passed through 2 mL of Ni-NTA nickel resin (Qiagen, 30250) pre-equilibrated with wash buffer (20 mM Tris-HCl pH 8.0, 300 mM NaCl, 30 mM imidazole). The resin was washed twice with 10 column volumes (CV) of wash buffer, and then eluted with 3 CV of elution buffer (20 mM Tris-HCl pH 8.0, 300 mM NaCl, 300 mM imidazole). The eluted proteins were concentrated using Ultra-15 Centrifugal Filter Units (Amicon) and further purified by using a Superdex 75 Increase 10/300 GL (GE Healthcare) size exclusion column in TBS (25 mM Tris-HCl pH 8.0, 150 mM NaCl). Fractions containing monomeric protein were pooled, concentrated, and snap-frozen in liquid nitrogen and stored at −80°C.

### Plasmid construction for RBD

The SARS-CoV-2 RBD (BEI NR-52422) construct was synthesized by GenScript into pcDNA3.1-with an N-terminal mu-phosphatase signal peptide and a C-terminal octa-histidine tag (GHHHHHHHH). The boundaries of the construct are N-328RFPN331 and 528KKST531-C^21^.

### Transient transfection

RBD proteins were produced in Expi293F cells grown in suspension using Expi293F expression medium (Life Technologies) at 33C, 70% humidity, 8% CO2 rotating at 150 rpm. The cultures were transfected using PEI-MAX (Polyscience) with cells grown to a density of 3.0 million cells per mL and cultivated for 3 days. Supernatants were clarified by centrifugation (5 min at 4000 rcf), addition of PDADMAC solution to a final concentration of 0.0375% (Sigma Aldrich, #409014), and a second spin (5 min at 4000 rcf).

### Purification of RBD

His tagged RBD was purified from clarified supernatants via a batch bind method where each clarified supernatant was supplemented with 1 M Tris-HCl pH 8.0 to a final concentration of 45 mM and 5 M NaCl to a final concentration of 310 mM. Talon cobalt affinity resin (Takara) was added to the treated supernatants and allowed to incubate for 15 min with gentle shaking. Resin was collected using vacuum filtration with a 0.2 mm filter and transferred to a gravity column. The resin was washed with 20 mM Tris pH 8.0, 300 mM NaCl, and the protein was eluted with 3 column volumes of 20 mM Tris pH 8.0, 300 mM NaCl, 300 mM imidazole. The batch bind process was then repeated and the first and second elutions combined. SDS-PAGE was used to assess purity. Following IMAC purification, the elution was concentrated and applied to a Cytiva S200 Increase column equilibrated with 20mM Tris 150mM NaCl pH8.0, and the peak of interest was collected and quantified using A280. The purified RBD was qualified using BLI to confirm binding using CR3022 and hACE2-Fc.

### In vitro bioluminescence characterization with monoclonal antibodies

A Synergy Neo2 Microplate Reader (BioTek) was used for all in vitro bioluminescence measurements. Assays were performed in 50% HBS-EP (GE Healthcare Life Sciences) plus 50% Nano-Glo assay buffer. For each well of a white opaque 96-well plate, 5 μL of 10× lucCage, 5 μL of 10× lucKey, 5 μL of 10× RBD, 5 μL of 10× antibody and remaining volume of buffer for in total 50 μL were mixed to reach the indicated concentration and ratio. The lucCage and lucKey components were incubated for 30 min at room temperature to enable pre-equilibration. The plate was centrifuged at 1*,000g* for 10 s and incubated at room temperature for a further 30 min. Then, 15 μL of 100× diluted furimazine (Nano-Glo luciferase assay reagent, Promega) was added to each well. Bioluminescence measurements in the absence of target were taken every 1 min (0.1 s integration and 10 s shaking during intervals) for a total of 30 min. To calculate the percent decrease in dynamic range for the graphs, the following formula was used:

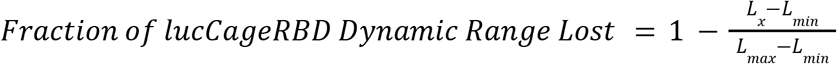

where L_x_ is the luminosity with 5 nM RBD and the tested antibody concentration, L_min_ is the luminosity when no RBD is added, and L_max_ is the luminosity when only 5 nM RBD is added. To derive half-maximal effective concentration (EC50) values from the bioluminescence-to-analyte plot, the top three peak bioluminescence intensities at individual analyte concentrations were averaged, subtracted from blank, and used to fit the sigmoidal 4PL curve.

### Detection of spiked RBD in human serum specimens

Serum specimens were derived from excess plasma or sera from adults (>18 years) of both genders provided by the Director of the Clinical Chemistry Division, the hospital of University Washington. All anonymized donor specimens were provided de-identified. Because the donors consented to have their excess specimens be used for other experimental studies, they could be transferred to our study without additional consent. All samples were passed through 0.22-μm filters before use. 5 μL of 10× monomeric RBD (10 or 1000 nM), 5 μL of 10× lucCage (10 nM), 5 μl of 10× lucKey (10 nM), 5 μl of 10× Antares2 (0.5 nM), and 5 μL of human donor serum or simulated nasal matrix were mixed with 1:1 HBS:Nano-Glo assay buffer to reach a total volume of 50 μl. The plate was centrifuged at 1*,000g* for 10 s. After 30 min incubation, 15 μL of 100× diluted furimazine in buffer was added to each well. Bioluminescence signals were recorded from both 470/40 nm and 590/35 nm channels every 1 min for a total of 30 min. The ratio *(R)* at each time point was calculated by the following equation as previously described^14^:

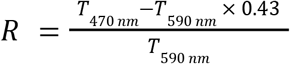

where T_470 nm_ and T_590 nm_ is the total luminescent signal at 470 nm and 590 nm, respectively. For calculating the fraction of lucCageRBD dynamic range lost for serum samples, the following equation was used:

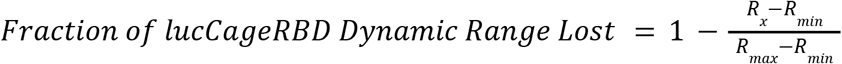

where *R_x_* is the *R* with 1 nM RBD in serum sample, *R_min_* is the *R* of serum sample but no RBD, and *R_max_* is the *R* of 100 nM RBD in the same serum sample. Monomeric SARS-CoV-2 RBD was expressed and purified as previously described^21^.

### In vitro bioluminescence characterization of lyophilized biosensors

5 μL of 10× lucCage and 5 μL of 10× lucKey were added to each well of a white opaque 96-well plate and lyophilized overnight. The biosensor was reconstituted in 10 μL of dH20 prior to testing, then 84 μL of 50% HBS-EP (GE Healthcare Life Sciences) plus 50% Nano-Glo assay buffer was added to each well. The lucCage and lucKey components were incubated for 30 min at room temperature to enable pre-equilibration. 1 μL of 100× diluted furimazine (Nano-Glo luciferase assay reagent, Promega) was added to each well. The plate was centrifuged at 1,000*g* for 10 s. Then, 5 μL of serially diluted target RBD were added to each well and measured in a A Synergy Neo2 Microplate Reader (BioTek). Measurements were taken every 1 min (0.1 s integration and 10 s shaking during intervals) for a total of 90 min.

### Biolayer interferometry

Protein-protein interactions were measured by using an Octet RED96 System (ForteBio) using streptavidin-coated biosensors (ForteBio). Each well contained 200 μL of solution, and the assay buffer was HBS-EP+ buffer (GE Healthcare Life Sciences, 10 mM HEPES pH 7.4, 150 mM NaCl, 3 mM EDTA, 0.05% (v/v) surfactant P20) plus 0.5% non-fat dry milk blotting grade blocker (BioRad). The biosensor tips were loaded with analyte peptide or protein at 20 μg mL^-1^ for 300 s (threshold of 0.8 nm response), incubated in HBS-EP+ buffer for 60 s to acquire the baseline measurement, dipped into the solution containing cage and/or key for 1800 s (association step) and dipped into the HBS-EP+ buffer for 1800 s (dissociation steps). The binding data were analysed with the ForteBio Data Analysis Software version 9.0.0.10.

### Live and pseudovirus entry and serum neutralization assays

SARS2-02 and SARS2-38 were assayed for neutralization potency by focus-reduction neutralization test (FRNT) as described previously^34^, and using Vero-TMPRSS2 cells. Briefly, serial dilutions of antibody were incubated with 2 x 10^2^ focus forming units of SARS-CoV-2 of the indicated strain for 1 h at 37°C in duplicate. Immune complexes were then added to Vero-TMRPSS2 cell monolayers in a 96-well plate and incubated for 1 h at 37°C prior to the addition of 1% (w/v) methylcellulose in MEM. Following incubation for 30 h at 37°C, cells were fixed with 4% paraformaldehyde (PFA), permeabilized and stained for infection foci with a mixture of mAbs that bind various epitopes on the RBD and NTD of spike (SARS2-02 and SARS2-38; diluted to 1 μg mL^-1^ total mAb concentration). Antibody-dose response curves were analyzed using non-linear regression analysis (with a variable slope) (GraphPad Software).

For the mAbs CV30, B38, and CR3022 and for the vaccinated human serum samples, HEK-hACE2 cells were cultured in DMEM with 10% FBS (Hyclone) and 1% PenStrep with 8% CO2 in a 37C incubator (ThermoFisher). Prior to plating, 40 uL of poly-lysine (Sigma) was placed into 96-well plates and incubated with rotation for 5 min. Poly-lysine was removed, plates were dried for 5 min then washed 1 × with water prior to plating 2×10^4^ cells. The following day, cells were checked to be at 80% confluence. In a half-area 96-well plate a 1:3 serial dilution of mAb or sera was made in DMEM starting at 1:10 initial dilution in 22 uL final volume. 22 uL of pseudovirus was then added to the serial dilution and incubated at room temperature for 30-60 min. HEK-hACE2 plate media was removed and 40 uL of the sera/virus mixture was added to the cells and incubated for 2 h at 37°C with 8% CO2. Following incubation, 40 uL 20% FBS and 2% PenStrep containing DMEM was added to the cells for 48 h. Following the 48-h infection, One-Glo-EX (Promega) was added to the cells in half culturing volume (40 uL added) and incubated in the dark for 5 min prior to reading on a Varioskan LUX plate reader (ThermoFisher). Measurements were done on all mAbs and human serum samples from each group in at least duplicate. Relative luciferase units were plotted and normalized in Prism (GraphPad) using a zero value of cells alone and a 100% value of 1:2 virus alone. Nonlinear regression of log(inhibitor) versus normalized response was used to determine IC50 values from curve fits.

### Sensor simulations

Mathematical models describing the ternary and quaternary sensor systems were simulated to test their responses to changes in their input parameters (concentrations and affinities of intervening species). Systems of ordinary differential equations describing the kinetics of the interactions between the species involved in each sensor were numerically integrated using integrate.odeint() as implemented in Scipy (version 1.6.3)^35^. Steady-state values were used to determine the distribution of species at thermodynamic equilibrium.

The ternary system is composed of the following species: ACE2, RBD, nAb, ACE2:RBD, RBD:nAb. The following set of equations was used to describe the system:

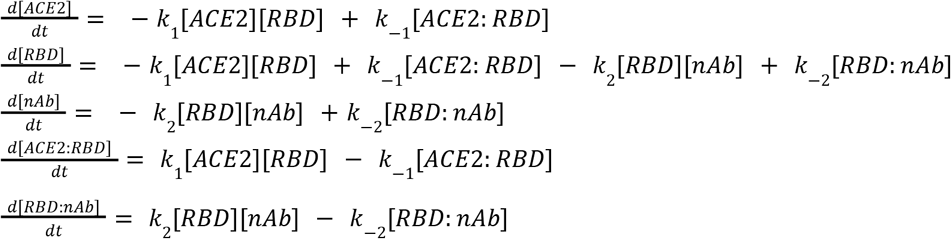

where *k_i_* describe bimolecular association rate constants and *k_-i_* represent unimolecular dissociation rate constants. *K*_1_ = *k*_-1_ / *k*_1_, and *K*_2_ = *k*_-2_ / *k*_2_ describe the equilibrium dissociation constants for the ACE2:RBD and RBD:nAb complexes respectively. For all ternary system simulations, *K*_1_ was set to 15 nM^36^. For consistency with the metric used to report on the quaternary system response (fraction of lucCageRBD dynamic range lost; described in section “In vitro bioluminescence characterization with monoclonal antibodies”), simulations for the the ternary system were reported as:

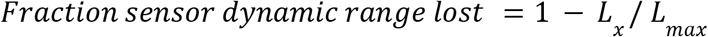

where *L_x_* is the signal observed when all three species are present, and *L_max_* is the response when nAb is absent.

The quaternary system is composed of the following species: cCL, oCL, lucKey, RBD, nAb, lucKey:oCL, oCL:RBD, lucKey:oCL:RBD, RBD:nAb. Only the open state of the Cage-Latch (oCL) is considered binding-competent, while the closed state (cCL) is not. The following set of equations was used to describe the system:

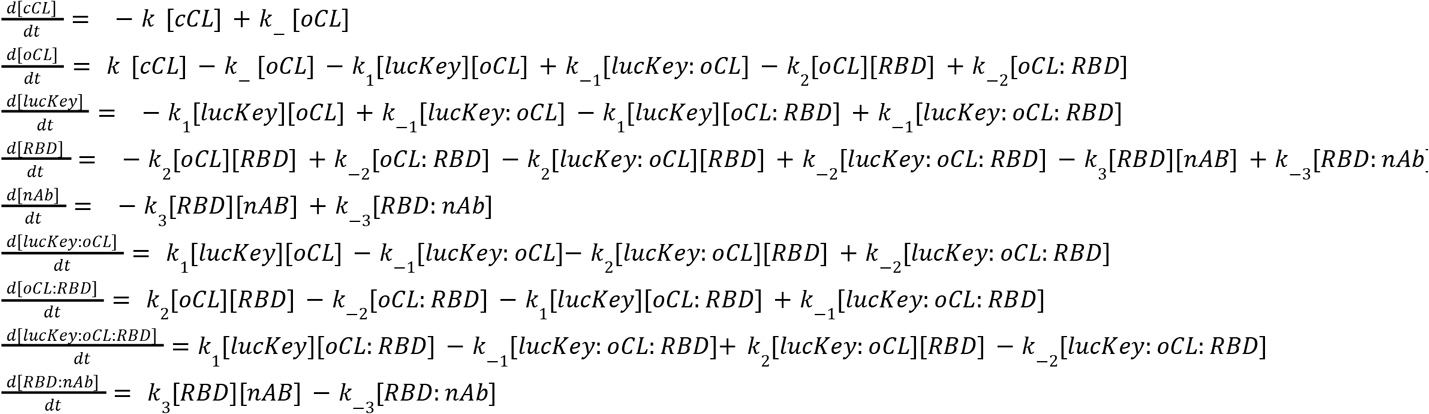

where *k_i_* describe bimolecular association rate constants and *k_-i_* represent unimolecular dissociation rate constants. *K*_1_ = *k*_-1_ / *k*_1_, *K*_2_ = *k*_-2_ / *k*_2_, and *K*_3_ = *k*_-3_ / *k*_3_ describe the affinities (equilibrium dissociation constants) for the binding interfaces lucKey:oCL, oCL:RBD and RBD:nAb respectively. The binding of lucKey and RBD to the open Cage-Latch species is considered symmetrical, *i.e.* non-cooperative, meaning that the binding events are independent. *K* = *k*_ / *k* = *exp*(Δ*G_open_* / *RT*) describes the unimolecular binding equilibrium of the Latch to the Cage, with Δ*G_open_* the free energy of Latch opening, *R* the universal gas constant, and *T* the thermodynamic temperature (set to 298.15 K for all simulations).

These systems were simulated over a range of species concentrations, as well as RBD:nAb affinities, to explore the behavior of each sensor, and gain insights into the influence of different variables on the position of the detection thresholds. The python code for running these simulations is provided as a Jupyter notebook: https://github.com/bwicky/covid_nAb_sensor_simulation.

### Statistical analysis

No statistical methods were used to pre-determine the sample size. No sample was excluded from data analysis, and no blinding was used. De-identified clinical serum samples were randomly used for spiking in target proteins. Results were successfully reproduced using different batches of pure proteins on different days. Unless otherwise indicated, data are shown as mean ± s.e.m., and error bars in figures represent s.e.m. of technical triplicate. Biolayer interferometry data were analyzed using ForteBio Data Analysis Software version 9.0.0.10. All data were analyzed and plotted using GraphPad Prism 8.

## Data Availability

The data that support the findings of this study are available from the corresponding author upon reasonable request. Source data are provided with this paper.

## Code Availability

The code for the sensor simulations is made available as an interactive Jupyter notebook at https://github.com/bwicky/covid_nAb_sensor_simulation.

## Acknowledgements

We acknowledge funding from the Henrietta and Aubrey Davis Endowed Professorship in the UW Department of Biochemistry (D.B.), the United World Antiviral Research Network (UWARN) one of the Centers Researching Emerging Infectious Diseases “CREIDs”, NIAID 1 U01 AI151698-01 (D.B., L.S., and H-.W.Y.), the Audacious Project at the Institute for Protein Design (D.B., H-.W.Y., W.Y.), Eric and Wendy Schmidt by recommendation of the Schmidt Futures (H-.W.Y. and A.Q-.R.), The Open Philanthropy Project Improving Protein Design Fund (D.B. and S.E.B.). Bill & Melinda Gates Foundation (OPP1156262 D.V.), the European Molecular Biology Organization (fellowship ALTF 139-2018 to B.I.M.W.), Gree Real Estate (D.B.), the National Institute of General Medical Sciences (R01GM120553 to D.V.), the National Institute of Allergy and Infectious Diseases (DP1AI158186 and HHSN272201700059C to D.V.), a Pew Biomedical Scholars Award (D.V.), Investigators in the Pathogenesis of Infectious Disease Awards from the Burroughs Wellcome Fund (D.V.), and Fast Grants (D.V.). This study also was supported by NIH grant R01 AI157155 (M.S.D.).

We also thank Robert Waterson and Wesley C. Van Voorhis for advice and support with the anti-SARS-CoV-2 antibody sensors and Brooke Fiala and Rashmi Ravichandran at the Institute for Protein Design for providing SARS-CoV-2 RBD and LCB1.

## Author contributions

D.B. conceived and supervised the project. H.-W.Y. and J.Z.Z. designed the research and performed all biosensor assays. D.V. supervised and A.C.W. and K.S. performed pseudovirus neutralization assay. B.I.M.W. created the thermodynamic equilibrium simulations of sensor systems. J.Z.Z., H.-W.Y., analyzed lucCageRBD and BLI data. L.A.V. and M.S.D. provided SARS2-02 and SARS2-38 antibodies and performed live virus neutralization experiments. R.T. and M.W. provided vaccinated human serum samples and performed ELISAs. M.N.P. and J.C.K provided vaccinated mouse serum samples and performed ELISAs. A.Q.-R. designed the lucCageRBD sensor and performed lyophilization experiments. M.D., C.C. and L.C. performed production and purification of proteins. J.Z.Z., H.-W.Y., B.I.M.W., L.S. and D.B. wrote the original draft. All authors reviewed and accepted the manuscript.

---

**Extended Data Fig 1:**
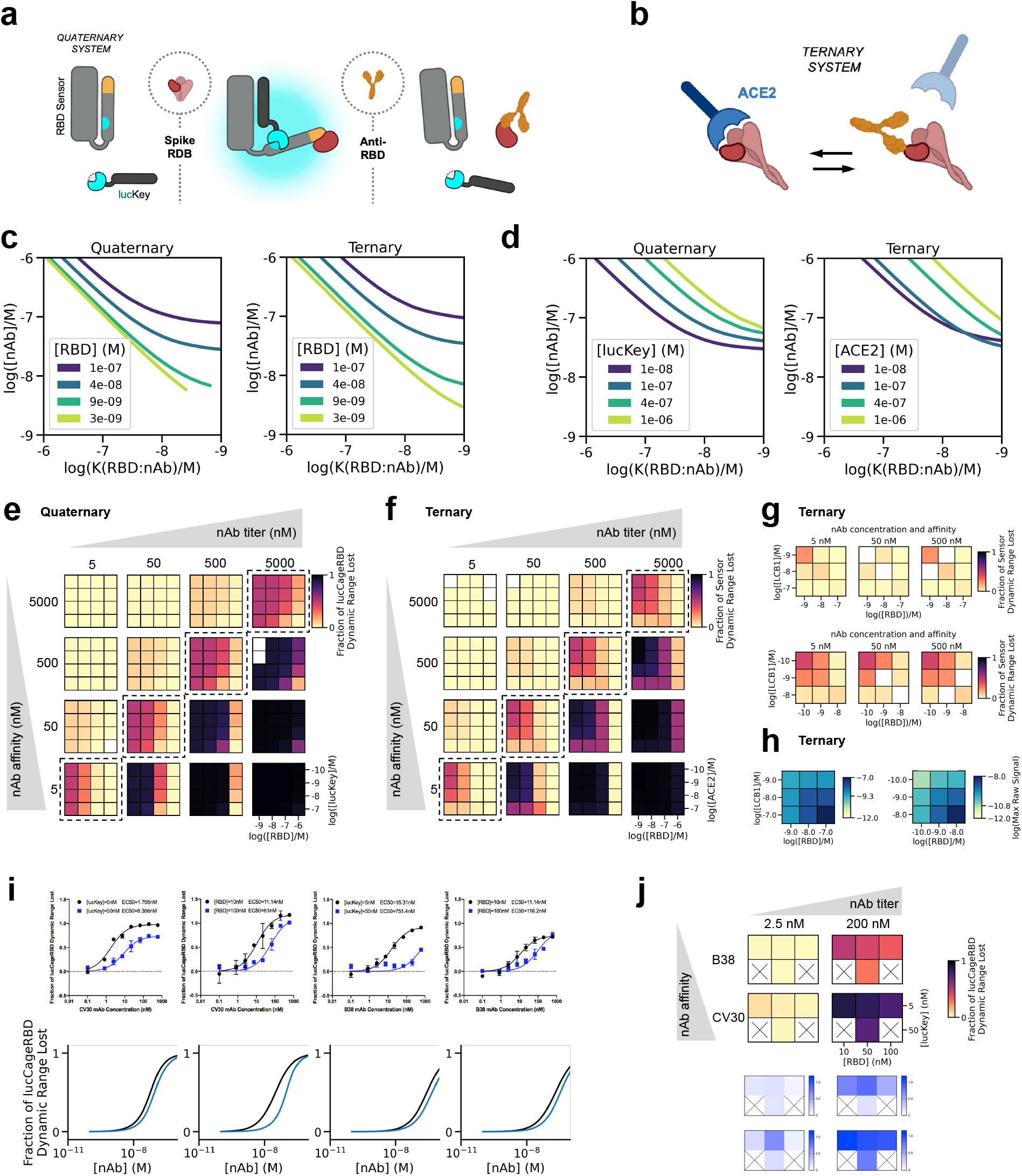
Simulation of biosensor systems. **a,** Schematic of the LOCKR nAb biosensor system (quaternary, this work), and **b**, ACE2:RBD out-competition format (ternary, previous work). **c-d**, Detection capabilities as a function of sensor settings (concentration of sensor components) along the dimension of each independent variable. The decision boundaries (EC50) are shown as solid lines. **c**, The concentration of RBD primarily affects the antibody titer detection limit. For tight antibodies (right-hand side of the plots), the position of the decision boundary (EC50) is solely determined by the concentration of RBD. The concentrations of the untitrated components were: Cage-Latch = 1 nM, lucKey = 50 nM for the quaternary system, and ACE2 = 1 nM for the ternary system. **d**, Effect of ACE2 and lucKey concentrations on the ternary and quaternary sensors respectively. Changing lucKey concentration in the quaternary case allows for a better modulation of the nAb affinity detection threshold. The concentration of RBD was set to 50 nM in both cases, and the concentration of Cage-Latch was set to 1 nM for the quaternary system. **e-f,** Decision matrices for the quaternary (**e**) and ternary (**f**) sensors. Each sub-matrix represents the simulated response for 16 different sensor settings (concentrations of sensor species, indicated at the bottom-right) for a given combination of nAb concentration and affinity. The quaternary system is capable of deconvoluting affinity and concentration across all combinations that activate the sensor (distinct patterns), while the ternary system returns the same reading for the cases where nAb affinity and concentrations are in the same range (diagonal, highlighted by dashed squares). **g**, Diagonal of decision matrices for ternary systems composed of RBD and LCB1 instead of ACE2. Sensor readings using the same concentrations of species as the ACE2:RBD system (top) or adjusted concentrations (bottom). **h**, Maximum raw signals for the ternary system sensor as in (**g**) with unmodified settings (left), and adjusted concentrations (right). The unmodified settings reduce the raw maximum signal range, but also reduce the detection capabilities of the sensor. Adjusting the settings to improve detection also increases the raw maximum signal range. **i**, Different concentrations of either CV30 (high affinity; *K*d = 25 nM) or B38 (low affinity; *K*d = 192 nM) mAbs with 1 nM RBD sensor and different concentrations of WT RBD and lucKey in the lucCageRBD assay (experimental and simulation data, top and bottom respectively). **j,** Heat map representation summarizing some of the experimental data from (**i**) for low and high concentrations of CV30 and B38 antibodies (simulation and experimental data, top and bottom respectively). All lucCageRBD experiments were performed in triplicate, representative data are shown, and data are mean ± s.e.m. The quaternary system was simulated with the following parameters: Δ*G_open_* = 4 *kcal/mol*, [*Cage – Latch*] = 1 *nM, K*(*lucKey: oCL*) = 5 *nM, K*(*oCL: RBD*) = 500 *pM*. The ternary system was simulated with *K*(*ACE*2: *RBD*) = 15 *nM,* except for **g-h**, where *K*(*LCB*1: *RBD*) = 500 *pM* was used instead. Cases where simulations did not converge are shown as white squares. All lucCageRBD experiments were performed in triplicate and data are mean ± s.e.m.

**Extended Data Fig 2:**
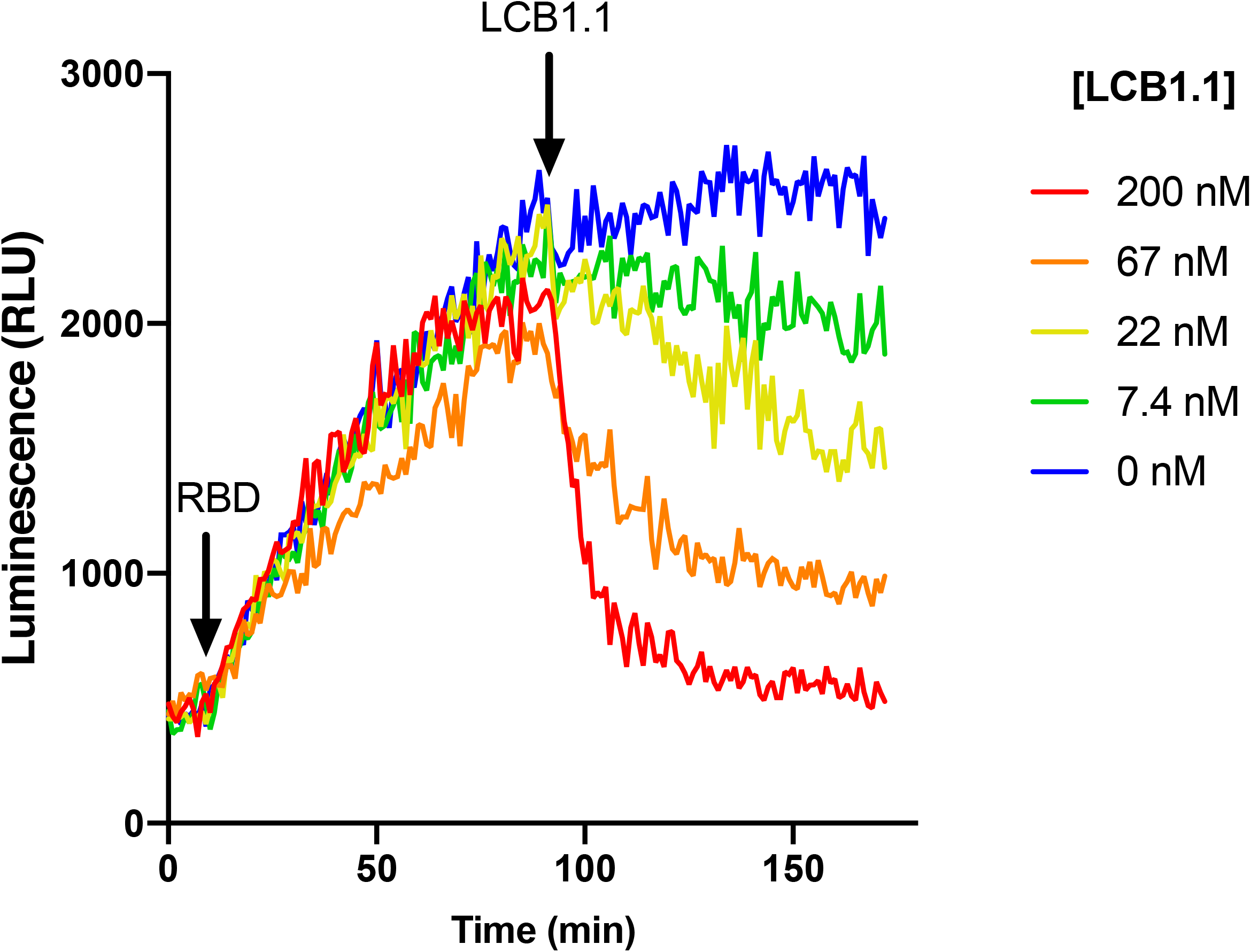
Reversibility of lucCageRBD. Time-course luminescence of the lucCageRBD assay using 200 nM of RBD WT and different concentrations of the *de novo* LCB1.1 binder^15^. All experiments were performed in triplicate, representative data are shown.

**Extended Data Fig 3:**
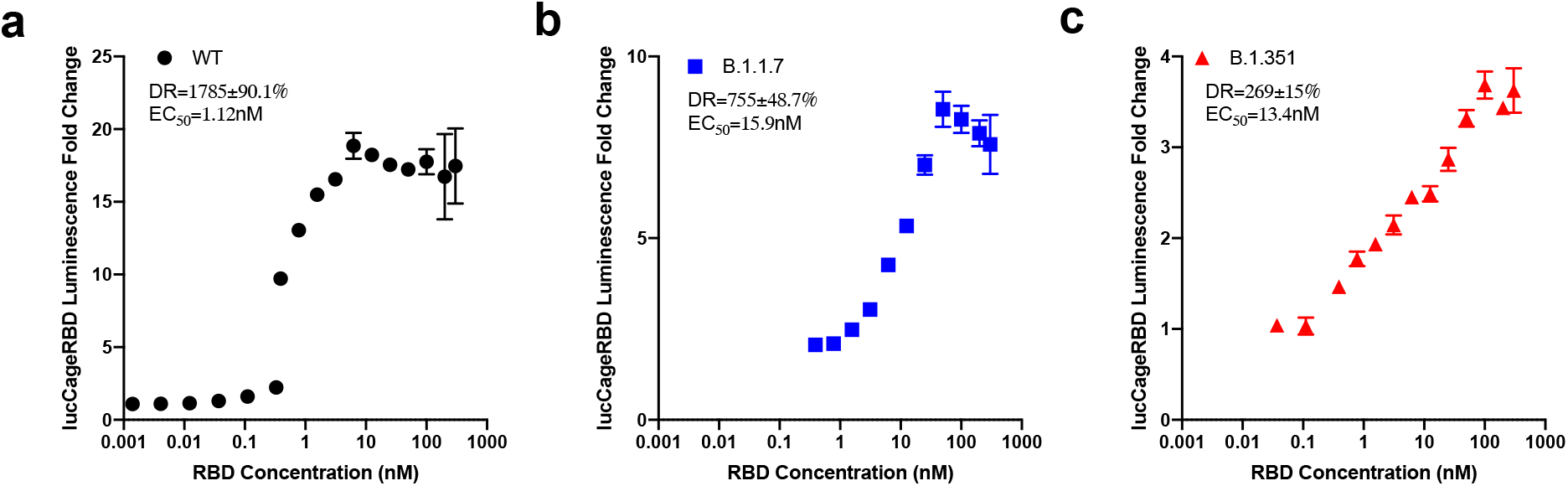
Detection of RBD strains by lucCageRBD. **a-c**, Fold change increase in luminescence from different concentrations of RBD WT (**a**), B.1.1.7 (**b**), and B.1.351 (**c**) tested in the lucCageRBD assay. All experiments were performed in triplicates and data are mean ± s.e.m.

**Extended Data Fig 4:**
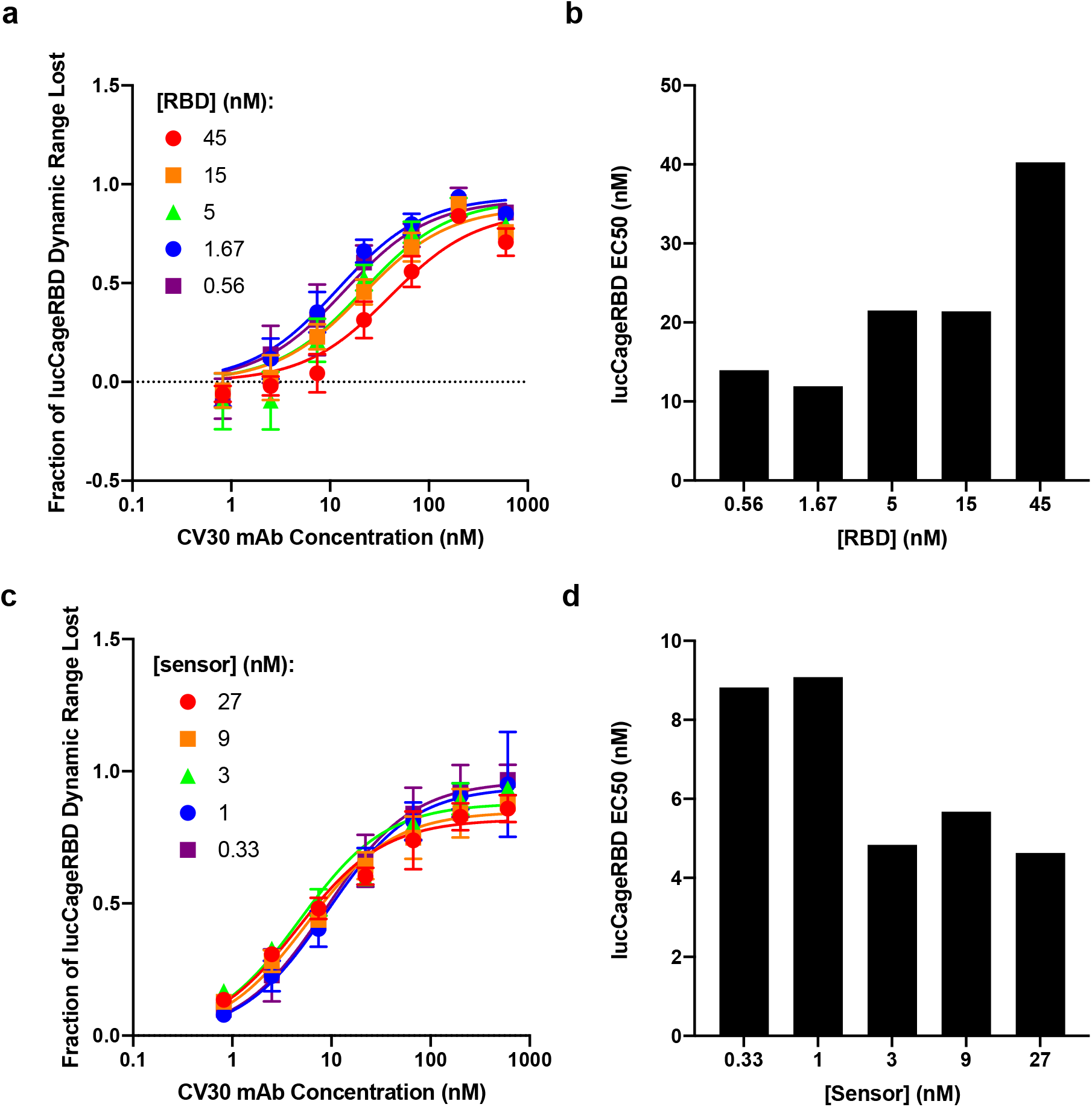
Effects of RBD and sensor titration on detecting SARS-CoV-2 antibodies. **a-b**, Resulting fraction of lucCageRBD dynamic range lost (**a**) and lucCageRBD EC50 (**b**) from different concentrations of mAb CV30 with different concentrations of RBD WT in 1 nM of RBD sensor and lucKey. **c-d**, Resulting fraction of lucCageRBD dynamic range lost (**c**) and lucCageRBD EC50 (**d**) from different concentrations of mAb CV30 with different concentrations of RBD sensor and lucKey (1:1 stoichiometry maintained) and with 5 nM RBD. All experiments were performed in triplicates and data are mean ± s.e.m.

**Extended Data Fig 5:**
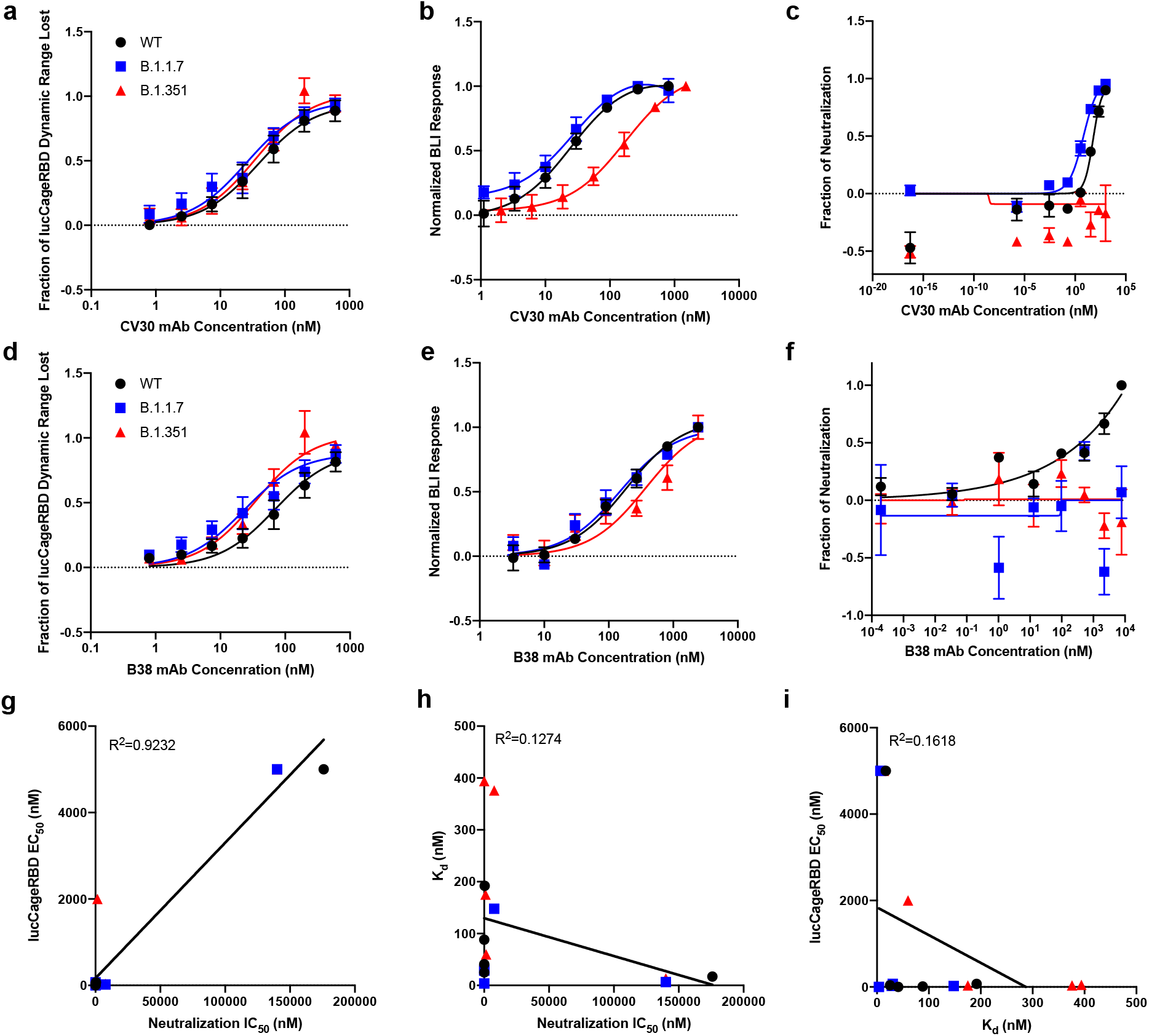
Monoclonal antibodies tested for lucCageRBD, binding, and neutralization. **a-f**, Different concentrations of either CV30 (**a-c**) or B38 (**e-g**) mAbs with 5 nM RBD WT, B.1.1.7, and B. 1.351 were tested in the lucCageRBD assay (**a, d**), BLI for binding to RBD strains (**b, e**), and spike VoC-presenting pseudovirus infection (VSV-based for **c**, HIV-based for **f**). Comparison of lucCageRBD EC50 and neutralization IC50 (**g**), binding affinity (k_d_) and neutralization IC50 (**h**), and lucCageRBD EC50 and k_d_ (**i**). All lucCageRBD and BLI experiments were performed in triplicate, neutralization experiments were performed in at least duplicate and data are mean ± s.e.m.

**Extended Data Fig 6:**
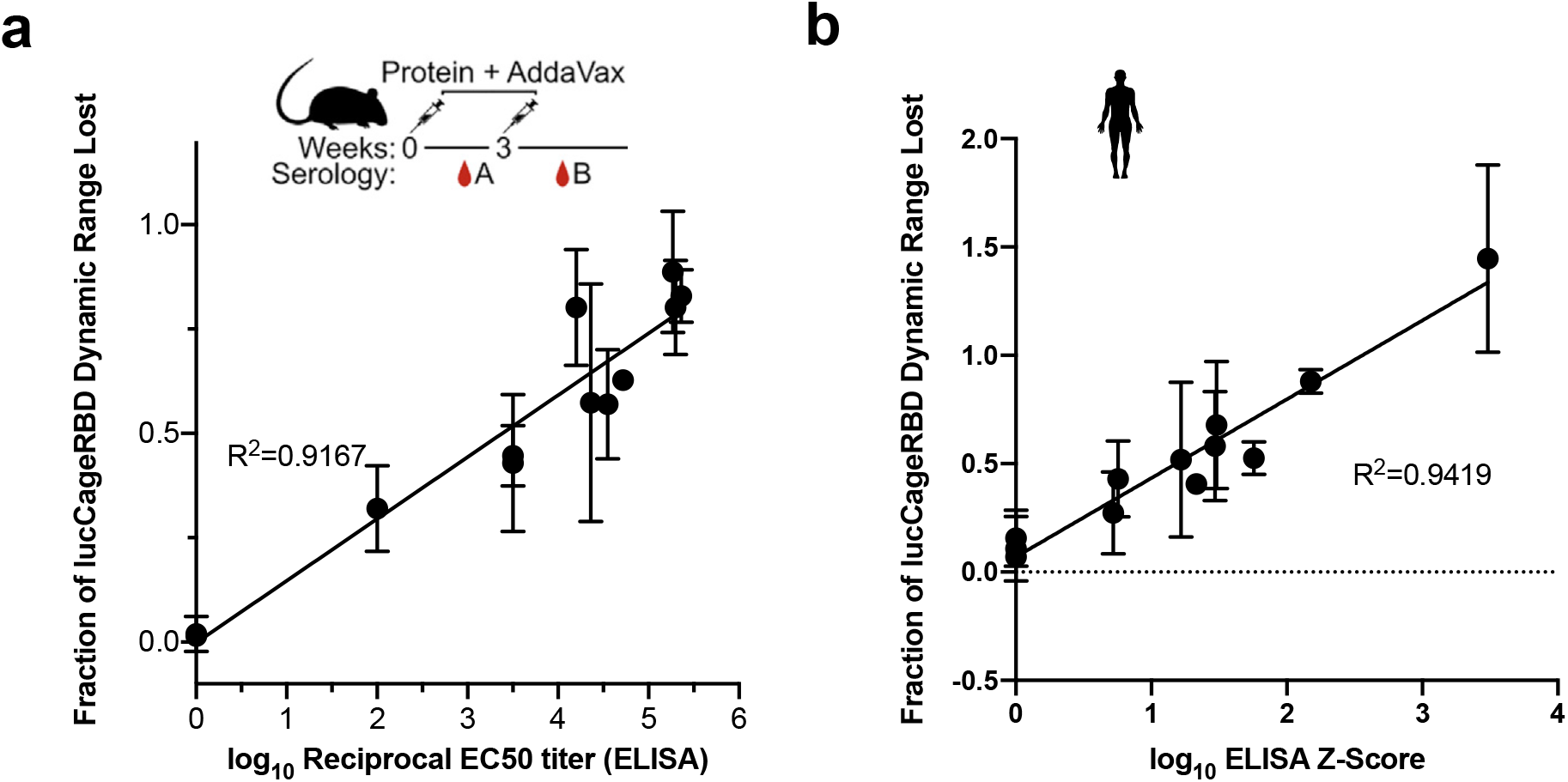
Detection of neutralizing antibodies in vaccinated serum. **a,** Serum (10%) from vaccinated mice were collected and tested for antibody concentration via ELISA (log_10_ reciprocal EC50 titer^37^) and in the lucCageRBD assay (n=12 serum samples). **b,** Serum (10%) from vaccinated patients were tested for antibody concentration via ELISA (log_10_ ELISA Z-score^23,33^) and in the lucCageRBD assay using RBD WT (n=12 serum samples). All experiments were performed in triplicates and data are mean ± s.e.m.

**Extended Data Fig 7:**
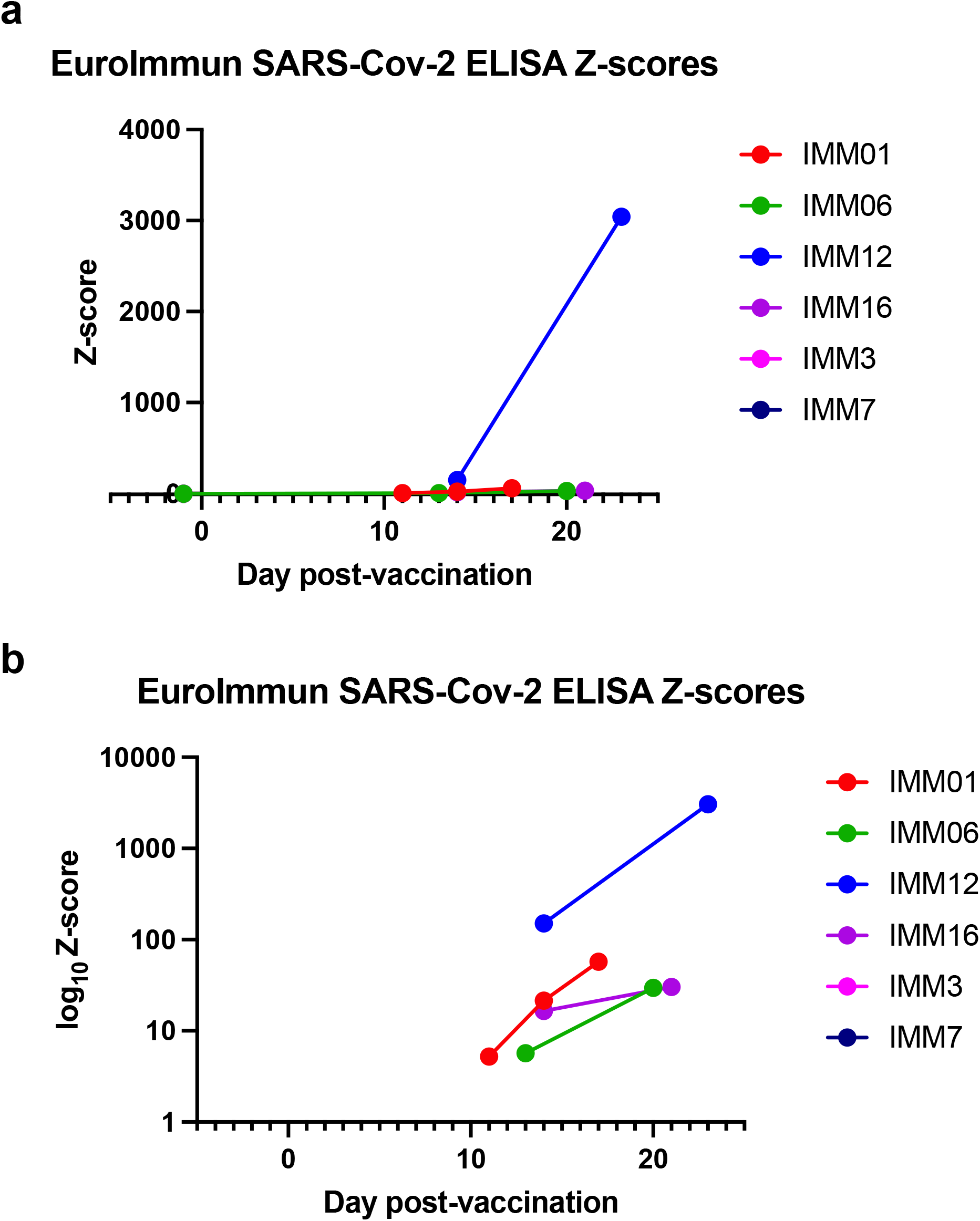
Antibody titer (ELISA) of vaccinated human serum. **a-b,** Serum collected from patients pre and post vaccination were tested in the EuroImmun ELISA for anti-spike antibodies. Patient’s anti-spike antibody levels (measured by Z-score^23,33^) were measured several days after vaccination. Z-score is plotted either in linear (**a**) or log_10_ scale (**b**). All experiments were performed in triplicate, representative data are shown.

**Extended Data Fig 8:**
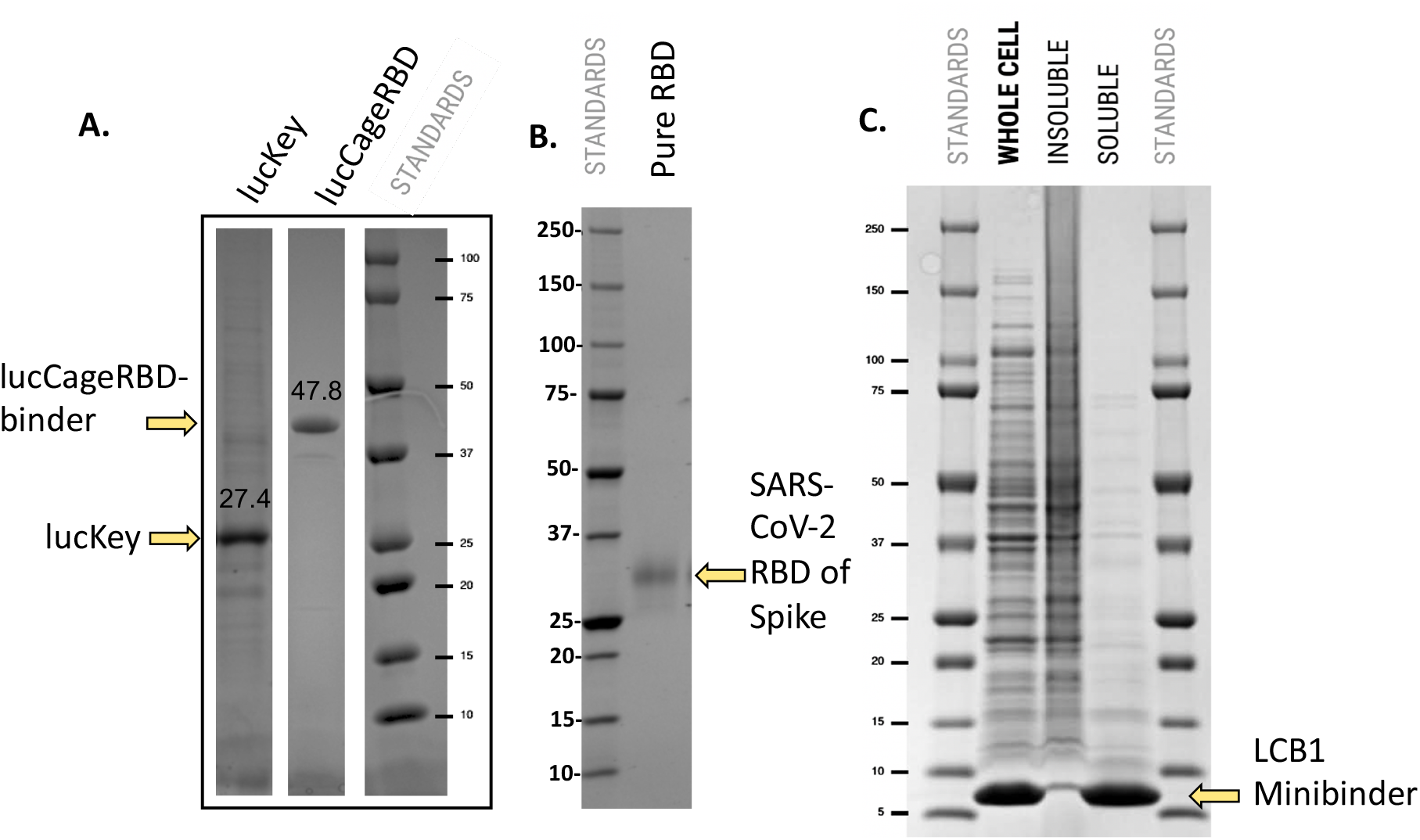
Production and purification of recombinant proteins used in LOCKR biosensor for serological diagnosis of neutralizing antibodies. **a,** Individual His6 tagged versions of LucKey (27.4 KDa) and lucCageRBD (47.8 KDa) are expressed in *E. coli* and following cell lysis are purified by IMAC using 50 mM imidazole in buffer elution. **b**, His6 tagged SARS-CoV-2 RBD (30 KDa) is expressed as a secreted protein in human 293 cells transiently transfected with DNA vectors encoding the protein which is purified from serum free culture medium by IMAC using 50 mM imidazole in buffer elution. **c**, His6 tagged LCB1 anti-SARS-CoV-2 RBD minibinder (7 KDa) is expressed in *E. coli* (whole cell). Following cell lysis by heating to 95°C and centrifugation to separate insoluble from soluble material, the LCB1 minibinder protein is >90% pure. Samples shown in A-C were analyzed by SDS-PAGE with reducing agent and Coomassie blue staining (molecular weight standards shown). All experiments were performed in triplicate, representative data are shown.

**Extended Data Fig 9:**
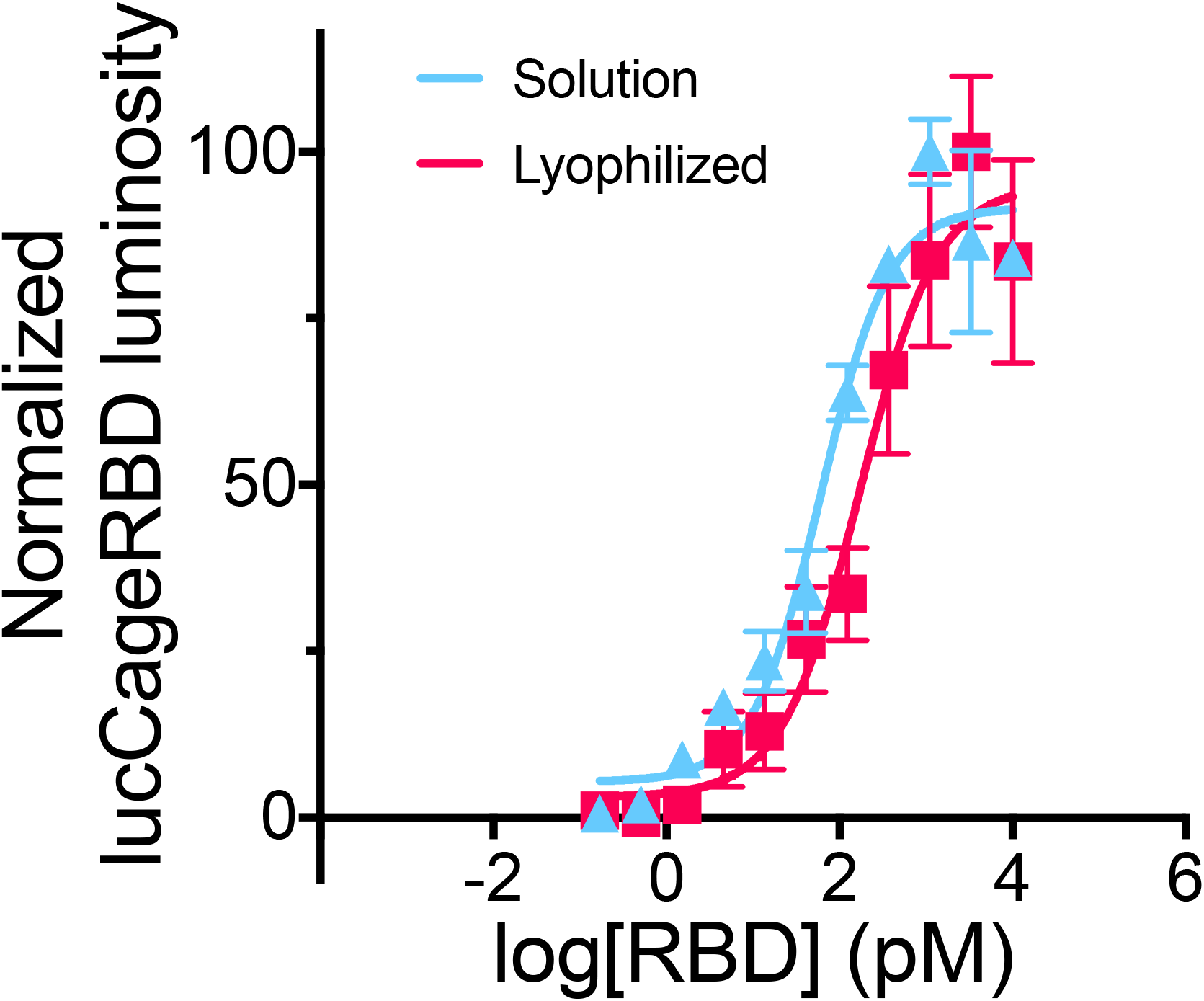
lucCageRBD biosensor sensitivity before and after lyophilization. lucCageRBD and lucKey were lyophilized at 10X concentration in a 96-well plate. After reconstitution in liquid format, the biosensor was tested at a final concentration of 1nM lucCageRBD and 1 nM lucKey for the detection of serially diluted RBD. All experiments were performed in triplicates and data are mean ± s.e.m.

